# UniSPAC: A Unified Segmentation Framework for Proofreading and Annotation in Connectomics

**DOI:** 10.1101/2024.11.27.625336

**Authors:** Juntao Deng, Jiamin Wu, Changmin Chen, Qihao Zheng, Zhaoxiang Zhang, Jingpeng Wu, Wanli Ouyang, Chunfeng Song

## Abstract

Reconstructing dense neuronal connections from volume electron microscopy (vEM) images is a critical challenge in neuroscience, driving the development of various automatic neuron segmentation methods^1^. Although current state-of-the-art automated segmentation methods can achieve high segmentation accuracy, they still require substantial manual proofreading and rely heavily on labeled datasets, which are often scarce, particularly for non-model organisms. Here, we introduce a **Uni**fied **S**egmentation framework for **P**roofreading and **A**nnotation in **C**onnectomics (**UniSPAC**) by providing the interactive segmentation model in 2D-level and the neuron tracing model in 3D-level. UniSPAC-2D allows users to correct its segmentation errors through point-based prompts, combining segmentation and proofreading in a single framework. UniSPAC-3D automatically traces neurons segmented by UniSPAC-2D across image slices, significantly reducing human involvement. Furthermore, UniSPAC-2D and UniSPAC-3D models can facilitate the semi-automatic generation of labeled data for new species, eliminating the need for external annotation tools. The fresh annotated data generated during proofreading in turn optimizes the interactive model through an online learning strategy, reducing the labeling effort for novel species over time. UniSPAC outperforms the start-of-the-art Segment Anything Model (SAM) in Drosophila segmentation, achieving 47x higher efficiency, and surpasses ACRLSD in cross-species segmentation on zebra finch data.

## Introduction

Reconstructing the dense neuronal connections, known as connectomics, is critical for advancing our understanding of nervous system organization. Various imaging techniques, including magnetic resonance imaging (MRI), light microscopy, and volume electron microscopy (vEM), are being utilized to study connectomics across millimeter, micrometer, or nanometer scales^2^. In terms of dense neuron reconstruction with synapses, vEM stands out as the only practical approach due to its exceptional 3D imaging resolution^3–6^. Nevertheless, the high resolution of EM data corresponds to a massive data size, which can reach tens of terabytes (TB) or even petabytes (PB)^1^. This property poses a significant burden in terms of data storage and transmission costs, limiting its usage to few laboratories^7,8^. Most importantly, the fineness of the EM data furthermore leads to a very exhaustive labeling workload.

Manual labeling is the most primitive and reliable method but equally the most expensive and time-consuming^9^. Annotation tools for 3D EM data, such as webKnossos^10^, have been fueling connectomics research and producing sufficient labeled data for several model organisms. As annotated data has accumulated and machine learning technology has developed over recent decades, computational image segmentation approaches^11,12^ were introduced into the connectomics research^13^, which spawned the research of neuron segmentation. A large number of neuron segmentation methods based on affinity graph^13–21^ or foreground mask^22–25^have been proposed. These automatic segmentation algorithms have facilitated the development of more efficient annotation tools^26–28^ and significantly reduced the manual labeling burden in the connectomics research. Nevertheless, despite their advantages, automated segmentation algorithms are not flawless. Segmentation errors such as merging or splitting neuronal voxels in EM images still occur, necessitating follow-up proofreading^1,29,30^. For example, a recent report asserts that correcting the errors in the automatic reconstruction of a whole fly brain still required around 33 person-years of human proofreading^31^. Given the continued challenge of labor-intensive manual interventions, an artificial intelligence-based system called RoboEM was recently proposed as a solution to replace manual proofreading^32^. According to reports, the computational annotation cost for cortical connectomes using RoboEM was about 400-fold lower than manual error correction. However, the amount of proofreading required is closely related to the generalization performance of automatic segmentation algorithms, particularly across different brain regions and even species. For newly studied species, poor performance from pre-trained segmentation models can result in heavier proofreading workloads—sometimes even exceeding the effort of pure manual annotation. Therefore, the current paradigm for connectomics reconstruction of new species still involves multiple stages, including data annotation, automatic neuron segmentation model training, and error proofreading. Crucially, these processes are often carried out using disparate tools, placing additional strain on connectomics research.

Despite the progress made so far, current computational methods still face several challenges and limitations. First, the majority of existing research regards neuron segmentation and proofreading as two distinct processes using different tools, which is cumbersome. However, drawing upon the foundation model of semantic segmentation, SAM^33^, we posit that neuron segmentation and proofreading can be accomplished within a unified framework through the interactive segmentation functionality. Second, the current best-performing neuron segmentation methods^16,23^ were all developed based on deep learning, creating a strong dependency on massive and reliable annotated data. Although well-trained automatic segmentation methods could greatly reduce annotation workload, the initial manual labeling to acquire training data still remains indispensable and time-consuming, especially for those less studied non-model organisms^34,35^. Nevertheless, inspired by mEMbrain^26^, we suggest that data annotation and error proofreading can also be integrated into a unified framework, which could further accelerate the adaptation of segmentation models to new species data. Therefore, developing a unified framework that combines data annotation, segmentation, and proofreading in connectomics is both urgent and critical for advancing the field. Numerous studies leveraging SAM-based methodologies (e.g., microSAM^36^, TriSAM^37^ and webknossos^10^ integrated with SAM) are actively striving to develop such a unified framework for vEM image segmentation. However, despite their potential, SAM-based models exhibit limitations in efficiently handling multi-instance segmentation of densely packed vEM neurons. This inefficiency arises from their reliance on many precise prompts to accurately delineate the segmentation targets.

To address these challenges, we proposed a framework named UniSPAC for the 3D neuron segmentation in vEM images. We decompose the 3D neuron segmentation task into two key components: interactive 2D segmentation of neurons on individual slices and 3D neuron tracking across consecutive slices. To accommodate the segmentation and proofreading of neurons into the same framework, we first proposed the UniSPAC-2D. In contrast to conventional segmentation models, UniSPAC-2D was capable of understanding point prompts derived from human interaction. Therefore, the proofreader could correct the errors of UniSPAC-2D by modifying the point prompts, especially for notorious merge errors in neuron segmentation problems. Building on the segmentation results from UniSPAC-2D, we further introduced UniSPAC-3D to automatically track specified neurons across slices, thus remarkably reducing manual involvement. By incorporating both segmentation and proofreading into the 3D neuron segmentation process, UniSPAC-2D and UniSPAC-3D improve adaptability to vEM data from new species. Proofreading generated fresh annotated data, which in turn optimized the interactive model. Ultimately, driven by the loop of segmentation, proofreading, and online learning processes, the UniSPAC framework could attain a segmentation performance close to manual labeling. In terms of interactive segmentation, UniSPAC-2D demonstrates more powerful multi-neuron segmentation performance than SAM on Drosophila data, and also improves the interaction efficiency by 47 times. Besides, UniSPAC-3D further reduces the interaction workload exponentially. In terms of cross-species segmentation, UniSPAC demonstrates better generalization performance than ACRLSD on zebra finch data.

## Results

### Overview of the UniSPAC-2D and UniSPAC-3D models

Traditional paradigms for automatic neuron segmentation in vEM data typically rely on manually labeled data to train models specialized for specific species. It has been shown that with sufficient training data, current automatic segmentation methods are capable to guarantee excellent segmentation performance for test vEM data of the same species^16,23^. However, when a new species under investigation has emerged, existing automated segmentation models often struggle to achieve satisfactory accuracy. As shown in **Fig. 1a**, a segmentation model trained on manually labeled Drosophila data tends to produce numerous errors when used to segment zebra finch data and cannot correct these errors autonomously. Besides, proofreading after segmentation using tools like CAVE^29^ could be very time consuming since the neuronal vEM images of Drosophila and zebra finch have pronounced pattern differences. Inspired by the foundation model SAM^33^, we proposed a neuron segmentation paradigm that enables interactive correction of segmentation results through point prompts. While interactive models may not initially perform well on data from a new species, the interaction prompts can be refined iteratively until optimal segmentation results are achieved. These optimal results can then serve as newly labeled data and be used to fine-tune the interactive model’s parameters through online learning. With constant positive feedback, the interactive model could significantly boost the segmentation accuracy for new species through the continuous generation of annotated data.

**Fig 1.**
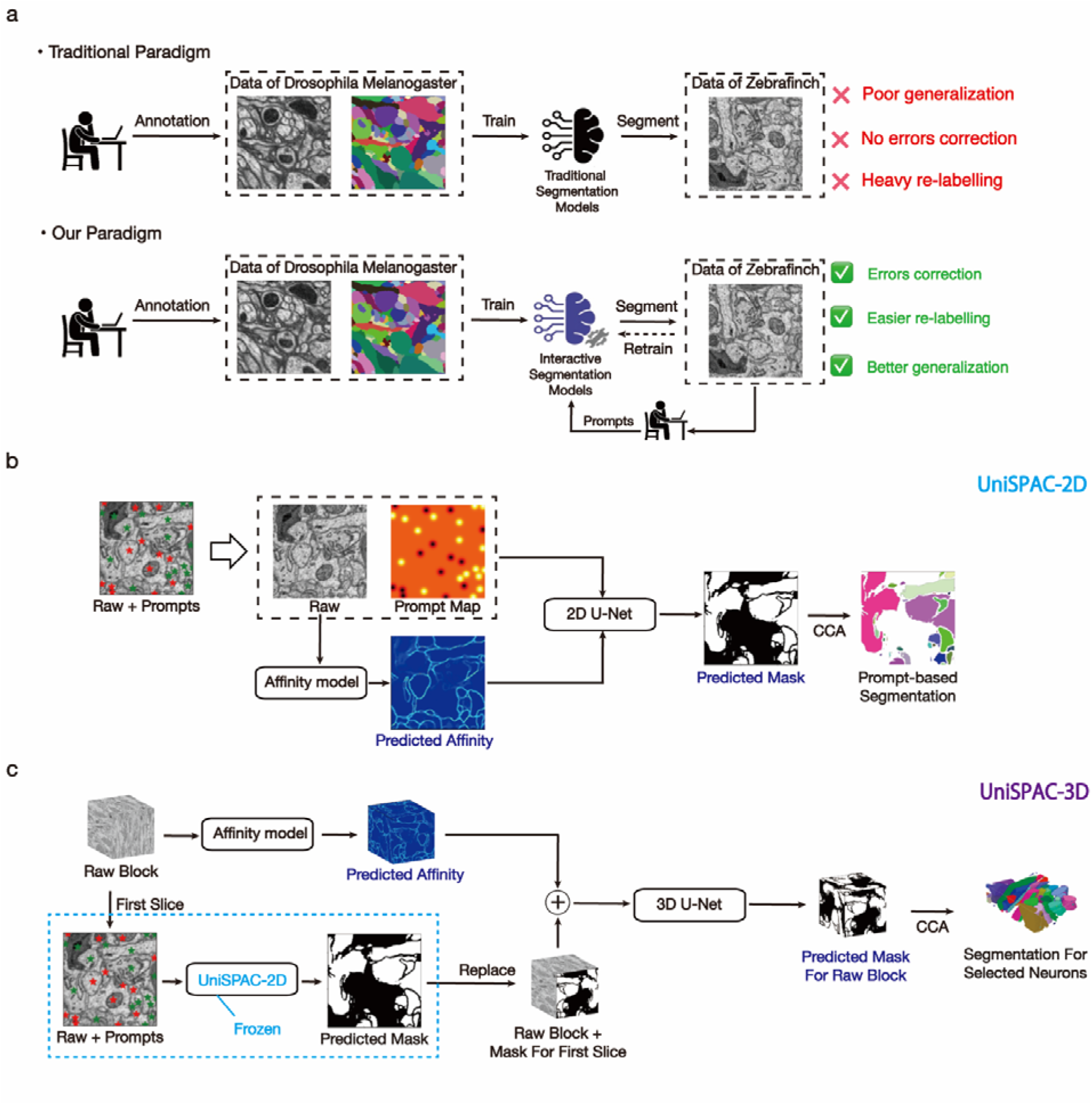
Overview of the UniSPAC-2D and UniSPAC-3D models. **a**, motivation and advantages of building the interactive neuron segmentation models. **b**, the model structure of UniSPAC-2D, which is adopted to segment neurons on the 2D slices. **c**, the model structure of UniSPAC-3D, which is employed to trace neurons across slices.

Due to the limitations of imaging techniques, three-dimensional data of electron microscopic neurons are typically constructed by stacking images of two-dimensional slices. Specifically, we split the task of interactive segmentation of 3D neurons into two subtasks: 1) interactive segmentation of neurons on 2D slices; 2) 3D tracking of selected neurons across slices. Based on this paradigm, we developed UniSPAC-2D (**Fig. 1b**) for 2D interactive segmentation and UniSPAC-3D (**Fig. 1c**) for 3D tracking. UniSPAC-2D is capable of predicting the masks of selected neurons based on user-supplied interaction information, which includes both positive and negative point prompts. Positive point prompts are used to indicate neurons to be segmented, while negative prompts mark regions to be excluded. The point prompts are converted into a graph format as detailed in Methods section. UniSPAC-3D then leverages the 2D masks generated by UniSPAC-2D as prompts to track neurons across slices, ultimately producing 3D segmentation masks of the selected neurons.

### Interactive segmentation of neurons for electron microscopy images

Existing neuron segmentation methods including FFN^23^ and ACRLSD^16^ are automatic segmentation algorithms that have demonstrated excellent performance in model organisms with sufficient data. However, they are still prone to errors, making manual proofreading an essential step. Currently, proofreading can only be finished with the help of additional proofreading tools like CAVE^29^ and RoboEM^32^, which adds complexity to the process. It would be far more convenient if both neuron segmentation and proofreading could be integrated into a single framework. The widely adopted interactive segmentation model, SAM^33^, provides a promising direction for uniting segmentation and proofreading. SAM accepts prompts in various formats (points, boxes, and text) and has shown remarkable generalization performance across a wide range of image segmentation tasks. Here, we offer a direct comparison between SAM and UniSPAC-2D for neuron segmentation, as SAM is currently limited to 2D image segmentation.

As demonstrated in **Fig. 2a**, we qualitatively compared the ability of SAM and UniSPAC-2D in segmenting different numbers of neurons. Both SAM and UniSPAC-2D provide good segmentation of individual neurons. However, as the number of neurons to be segmented increases, SAM begins to make errors. In the case of providing only the positive point prompt for each neuron, SAM tends to bias the entire image toward a foreground mask as the number of neurons grows. In contrast, UniSPAC-2D effectively handles the simultaneous segmentation of multiple neurons. The quantitative comparison results in **Fig. 2b** further demonstrate that UniSPAC-2D performs slightly better than SAM for single neuron segmentation, and its performance improvement in multi-neuron segmentation is significant. This highlights a limitation of SAM—it can only segment neurons one at a time when applied to vEM images (**Fig. 2c**). In contrast, UniSPAC-2D can support two modes when segmenting neurons: 1) automatically segment all neurons when the prompt information is absent, and 2) selectively segmentation when there exist prompts (**Fig. 2d**). In general, neurons are very dense in vEM images. For instance, in a 2µm^3^ training volume of HEMI-BRAIN, the number of neurons in each slice along the z-axis direction ranges from 80 to 120 (**Fig. 2e**). Besides, there are an average 95 neurons on each 2µm^.2^ slice. Thus, SAM would theoretically require about 95 interactions to fully segment a single slice. In contrast, UniSPAC-2D can complete the segmentation of one slice with just one automatic segmentation followed by two interactive segmentations (**Fig. 2f**). In summary, for a 2µm^3^ neuron vEM image of Drosophila, SAM requires approximately 47 times more interactions than UniSPAC-2D.

**Fig 2.**
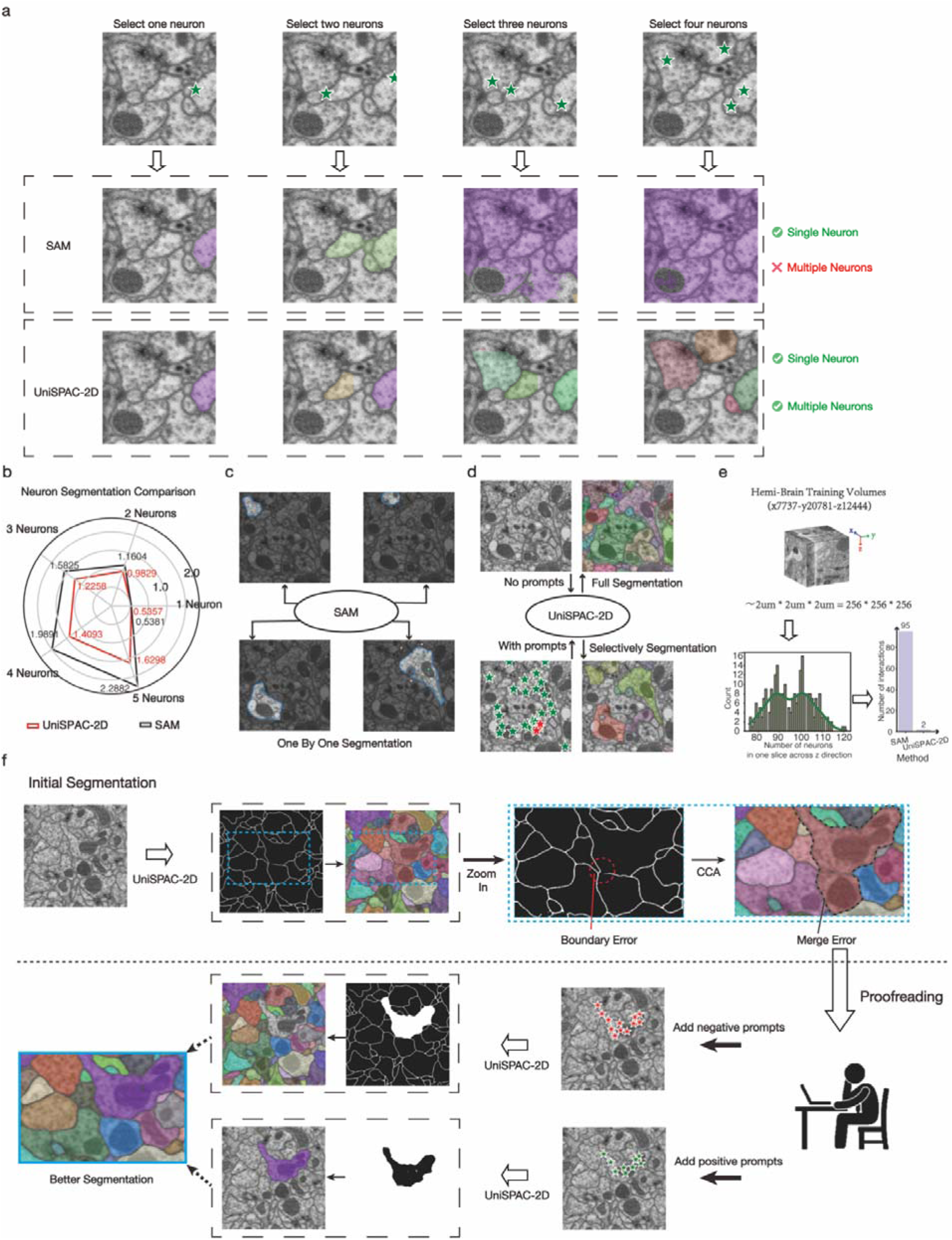
Performance demonstration of interactive neuron segmentation. **a**, Comparison of SAM and UniSPAC-2D at the same prompts. Columns 1 to 4 represent the number of neurons segmented in each slice, increasing from 1 to 4. A positive prompt was randomly assigned for each neuron. **b**, the quantitative comparison of the interactive segmentation performance of SAM and UniSPAC-2D. Segmentation performance was tested every ten slices on HEMI-BRAIN ROI-1, with the average Variation of Information (VOI) calculated over 100 tests. Lower VOI values indicate better segmentation performance. **c**, workflow for adopting SAM in segmentation of electron microscopy neuron images. **d**, workflow for employing UniSPAC-2D in segmentation of electron microscopy neuron images. **e**, statistics of the number of neurons per slice in Drosophila electron microscopy neuron data, along with a comparison of the number of interactions required for UniSPAC-2D and SAM. **f**, demonstration of the merge error correction process using UniSPAC-2D.

### Automatic tracking of 3D neurons

Although satisfactory segmentation results can be achieved using UniSPAC-2D, interactively segmenting vEM data slice by slice remains inefficient. To address this, we employed UniSPAC-3D to automatically segment neurons across slices. As illustrated in **Fig. 3a**, the 3D image block formed by stacking numerous slices was divided into several patches in the direction of the slices. For the first slice of each patch, we adopted UniSPAC-2D to acquire the satisfactory segmentation for neurons of interest with manually provided point prompts. The resulting mask of the initial slice could serve as an input for UniSPAC-3D, which then predict the masks of 3D neurons for the remaining slices within the same patch. The above process is carried out in all patches to generate the complete 3D masks for the target neurons. The number of slices in each patch could be set to 8, 16, 24, 32, and so on, with the corresponding labeling workload decreasing exponentially as UniSPAC-3D handles more slices automatically.

**Fig 3.**
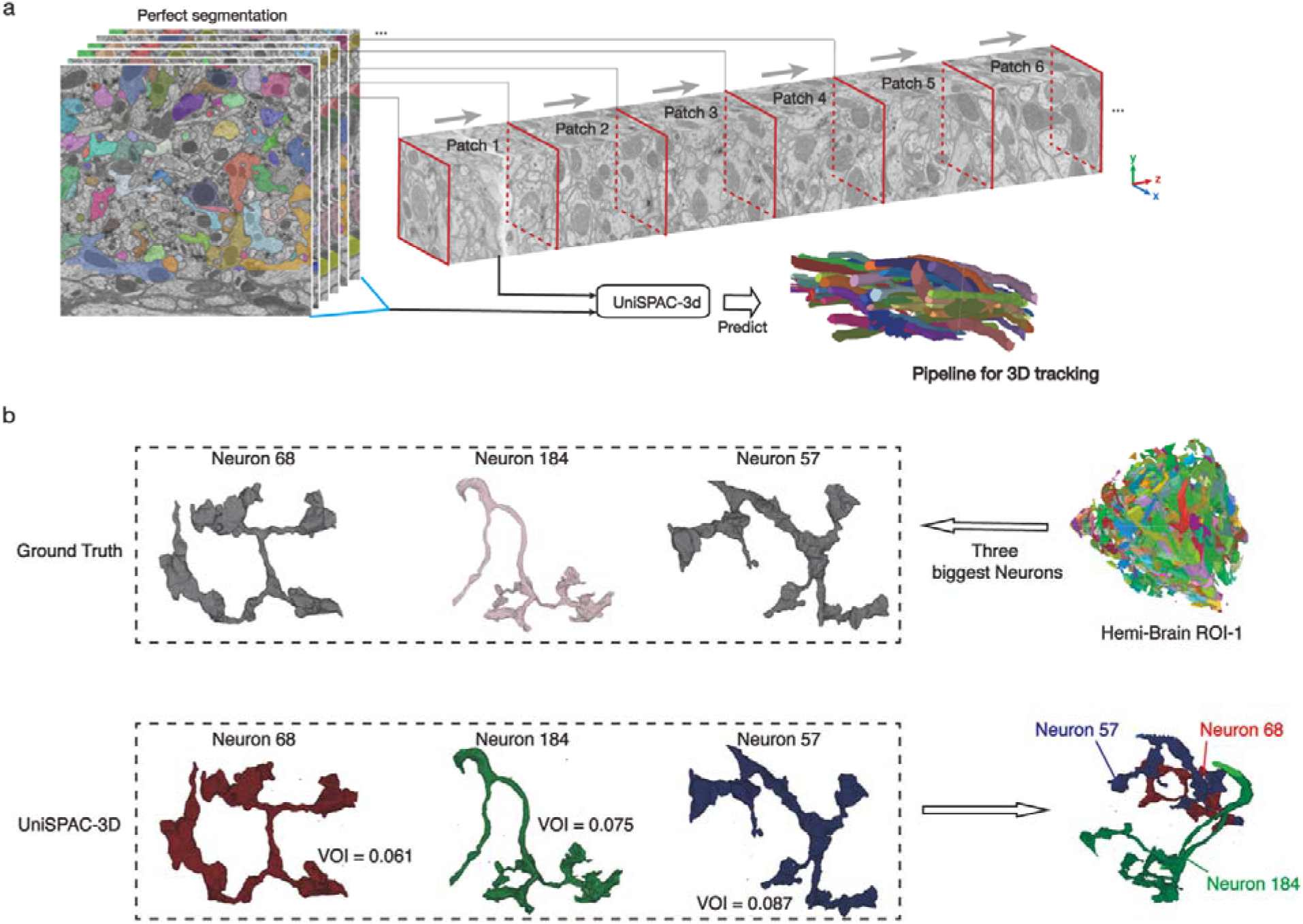
Process and results of neuron tracing. **a**, the procedure of tracking neurons specified by UniSPAC-2D across slices using UniSPAC-3D. **b**, The segmentation and tracing results for each of the three largest-volume neurons in HEMI-BRAIN ROI-1. The 3D visualizations of the neurons are shown along with the quantitative performance metric, Variation of Information (VOI).

We utilized 8 annotated volumes from HEMI-BRAIN and 4 annotated volumes from FIB-25 to train the neuron-tracking model UniSPAC-3D. In order to evaluate the segmentation accuracy of UniSPAC-3D, three neurons with the largest volume were selected from the HEMI-BRAIN ROI-1 (**Fig. 3b**). The entire HEMI-BRAIN ROI-1, which has a shape of ’1475×1475×1475,, was input into UniSPAC-3D for testing. However, only these three selected neurons are segmented and traced in this analysis. The operation should be conducted by employing UniSPAC-2D to obtain the optimal 2D masks of the three neurons on the initial slice of each patch, respectively. Here, we set the number of slices in each patch to 8 and tracked each neuron based on their 2D masks in the first slice using UniSPAC-3D. Considering that UniSPAC-2D is able to obtain optimal segmentation results through interaction, we directly employed the ground truth mask of the first slice of each patch as input in the inference of UniSPAC-3D. The results demonstrated that the variation of information (VOI) of all three neurons tracked by UniSPAC-3D were close to zero (**Fig. 3b**). It is worth noting that the VOI of the three largest neurons was also zero in the results of the FFN algorithm’s automated segmentation for the entire HEMI-BRAIN ROI-1. These results suggest that the tracking model UniSPAC-3D can achieve comparable segmentation performance to the start-of-the-art automated neuron reconstruction model FFN. Further experiments were conducted to test the impact of the patch size parameter on tracking performance (Supplementary Note 1).

### Interactive models speed up the annotation process for cross-species data

Existing automatic segmentation methods can deliver highly accurate segmentation results for species with sufficient manually labelled data. However, it should be noted that many species lack such manually labeled data. Given that traditional data annotation is time-consuming and labor-intensive, it is critical to use existing data more effectively to train robust cross-species segmentation models. Therefore, assessing the cross-species generalization capabilities of existing methods becomes essential. In this study, we adopted Drosophila melanogaster data to train the models and then tested them on zebra finch data. Three models were compared in this cross-species setting: ACRLSD-2D, UniSPAC-2D and UniSPAC-2D*. ACRLSD-2D is the 2D version of ACRLSD that we implemented to segment neurons at an agglomeration threshold of 0.5. UniSPAC-2D refers the version trained on 12 volumes of the Drosophila datasets (HEMI-BRAIN and FIB-25) with the combination of cross entropy loss and dice loss. Furthermore, UniSPAC-2D* is the enhanced version of UniSPAC-2D by incorporating a regularization term into the loss function during training. The augmented loss function resulted in UniSPAC-2D* exhibiting greater accuracy in predicting neuron boundaries. As shown in the test case of zebra finch in **Fig. 4a**, without any prompting, ACRLSD-2D achieved a VOI of 2.1093, while UniSPAC-2D and UniSPAC-2D* obtained VOIs of 2.2587 and 1.9752, respectively. Obviously, the neuron boundaries of in the mask predicted by UniSPAC-2D* are more continuous and coarser than those of UniSPAC-2D. This phenomenon can significantly reduce the merging errors generated by connected domain analysis. Nevertheless, there are still some zebra finch neurons with poorly predicted boundaries for ACRLSD-2D, UniSPAC-2D, and UniSPAC-2D*. As for these neurons with poor boundaries, it is nearly impossible for ACRLSD-2D to rectify these segmentation errors by itself. However, UniSPAC-2D* was able to predict the masks of these neurons with erroneous boundaries using point prompts provided by the proofreader (**Fig. 4a**). After combining two segmentation results, UniSPAC-2D* could achieve a VOI of 1.7734.

**Fig 4.**
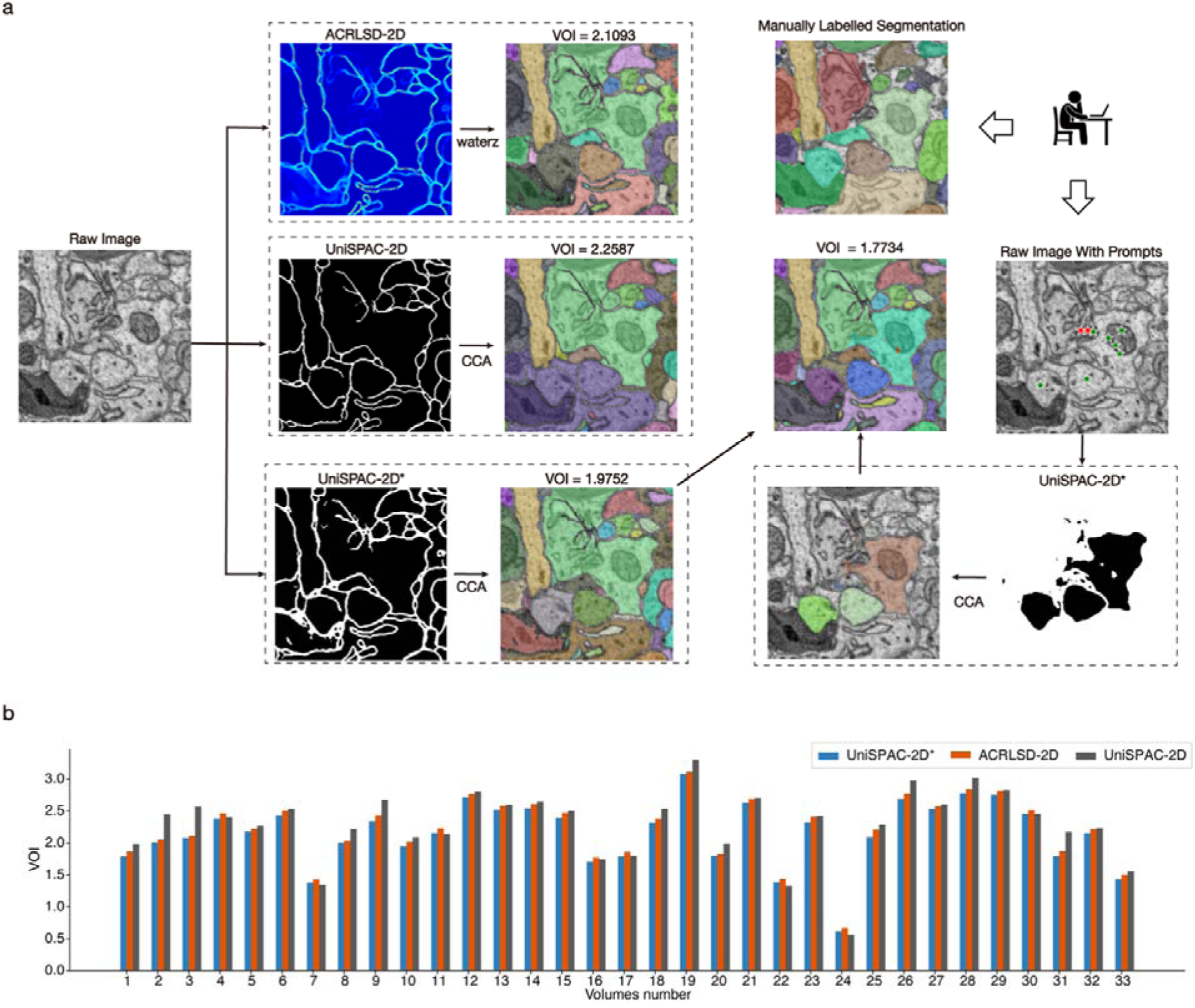
Performance comparison of different methods for cross-species neuron segmentation. Three models trained on Drosophila data were used for cross-species assessment: ACRLSD-2D, UniSPAC-2D and UniSPAC-2D*. ACRLSD-2D is the reproduced 2D version of the ACRLSD model. UniSPAC-2D was trained using a combination of cross entropy and dice loss. UniSPAC-2D* incorporates an additional loss term related to neuron boundaries to the original cross entropy and dice loss. **a**, Performance comparison of ACRLSD-2D, UniSPAC-2D and UniSPAC-2D* on a single slice from the zebra finch test data. **b**, quantitative comparison of the performance of the three models for zebra finch data across all 33 volumes. Both UniSPAC-2D and UniSPAC-2D* were set to the non-prompt mode.

To provide a more comprehensive comparison, the performance of ACRLSD-2D, UniSPAC-2D, and UniSPAC-2D* was evaluated on 33 volumes of zebra finch data. It should be noted that both UniSPAC-2D and UniSPAC-2D* made predictions without prompts here. As demonstrated in **Fig. 4b**, UniSPAC-2D achieved a better average VOI than ACRLSD-2D in only eight volumes, whereas UniSPAC-2D* achieved the best average VOI across all 33 volumes. These findings indicate that UniSPAC-2D* exhibits superior cross-species generalization performance in the non-prompt mode and performs even better when manual prompts are introduced. In addition, there is an elevation of VOI split (Supplementary Figure 3) for UniSPAC-2D* compared to UniSPAC-2D, while concurrently, a notable reduction in VOI merge has also been documented (Supplementary Figure 4).

### Online learning further improves segmentation performance on cross-species data

Given that the UniSPAC-2D model integrates segmentation and proofreading into a unified framework, it is feasible to employ this method for semi-automated annotation of new species data. The annotated data should be helpful in further optimizing the cross-species segmentation performance of UniSPAC-2D. In this study, we simulated the segmentation-proofreading-training pipeline using zebra finch data. Initially, UniSPAC-2D model was trained on 12 volumes of Drosophila data and then sequentially learned online over the front six zebra finch volumes. The remaining 27 zebra finch volumes were all adopted for testing. Seven different online learning strategies^38–42^ were employed and compared in terms of both the segmentation performance and time cost. The input prompts of UniSPAC-2D were set to the centroid of each neuron.

As shown in **Fig. 5a**, the VOI of UniSPAC-2D for zebra finch test data progressively decreased as more zebra finch volumes were used for training. Among the online learning strategies, the AGEM strategy^42^ achieved the closest performance to the baseline-1 strategy (training with all previously seen volumes). Besides, the AGEM strategy took only 58.2% of the training time of baseline-1 (**Fig. 5b**). Consequently, we ultimately chose the AGEM strategy for online learning. In **Fig. 5c**, we presented the variation of UniSPAC-2D’s segmentation results for one slice in zebra finch test data during online learning process. UniSPAC-2D model initially trained on the Drosophila data achieved poor segmentation results for the zebra finch slice in the case where each neuron only provided the centroid as prompts. The affinity graph predicted by the UniSPAC-2D model indicates that the poor segmentation performance is most likely due to the inability to predict neuron boundaries. However, as UniSPAC-2D was trained on more zebra finch volumes, its predicted affinity graph was significantly enhanced, raising neuron segmentation accuracy to match human labeling precision.

**Fig 5.**
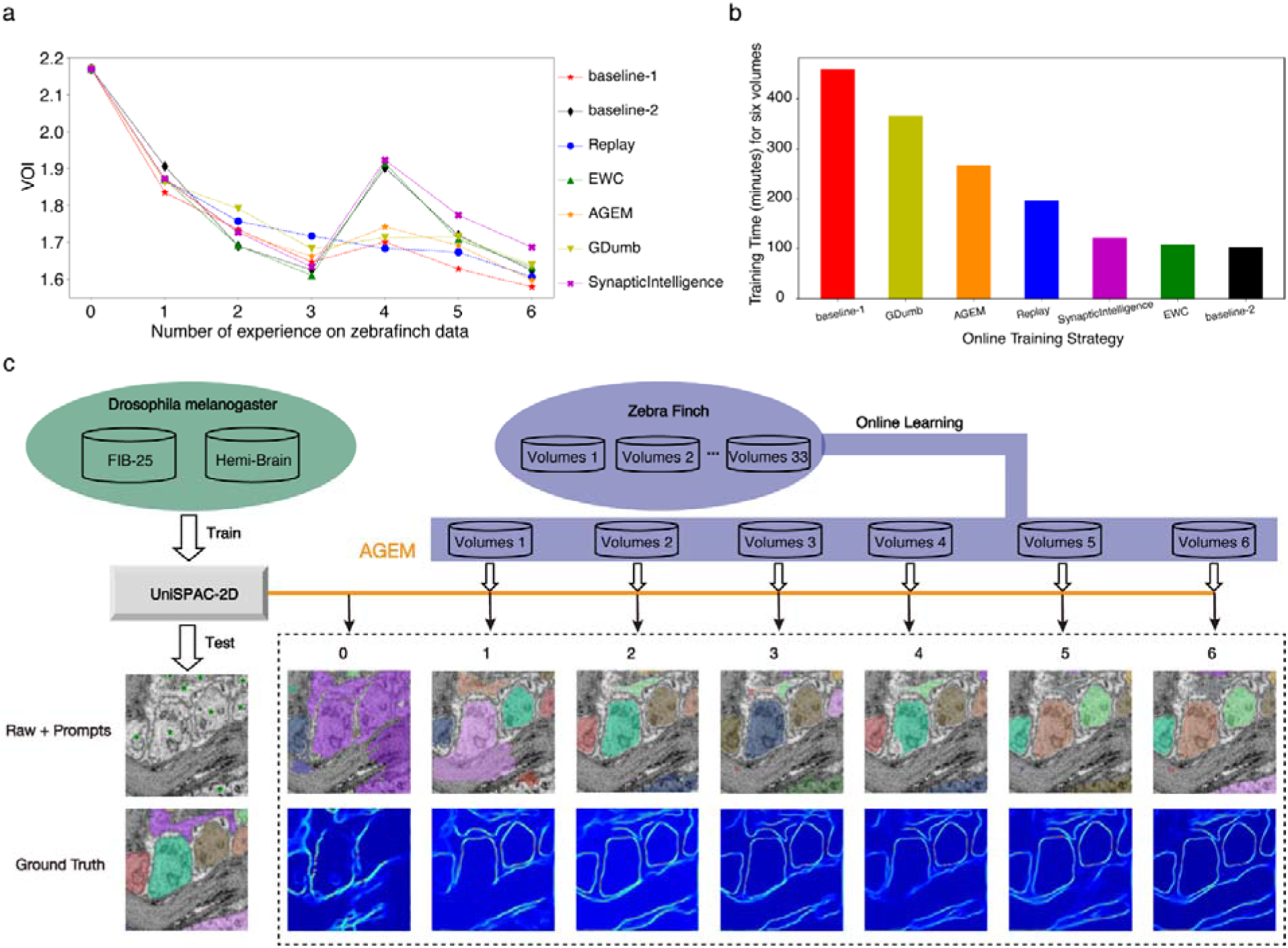
Online Learning Experiment Using Zebra Finch Data. Of the 33 zebra finch volumes, the first six were adopted as the training set for online learning, and the next 25 volumes were utilized for testing. **a**, comparison of the average performance of the UniSPAC-2D model on 25 test volumes using different online learning strategies. Baseline-1 refers to Cumulative Learning, where all previously seen zebra finch volumes are used in each training session. Baseline-2 involves fine-tuning UniSPAC-2D with only the current zebra finch volume in each training session. The initial UniSPAC-2D model was trained on Drosophila data. **b**, Time cost to train the UniSPAC-2D model with different strategies. **c**, visualization of performance improvement using the AGEM strategy on a test slice from the zebra finch volumes throughout the online learning process.

## Discussion

In summary, we proposed a paradigm for vEM neuron segmentation by introducing two models: the interactive segmentation model, UniSPAC-2D, and the neuron-tracking model, UniSPAC-3D. UniSPAC-2D can perform neuron segmentation and proofreading within the identical framework. As for the interactive segmentation in 2D vEM images of Drosophila neurons, UniSPAC-2D demonstrated performance far beyond the foundation model SAM. In addition, UniSPAC-2D makes the segmentation error correction easier, including the notorious merge error. Regarding segmenting vEM neurons across species, we introduced a loss function term that helped UniSPAC-2D acquire more continuous neuron boundaries in its output masks. On all 33 volumes of zebra finch, the enhanced UniSPAC-2D achieved smaller average VOIs compared to ACRLSD-2D. In addition, UniSPAC-2D could perform neuron proofreading even without prior exposure to zebra finch data, contributing to a better cross-species segmentation performance. Building on the segmentation results of UniSPAC-2D, UniSPAC-3D can trace the selected neurons across slices, which significantly reduces the amount of manual interaction. The tracking results for the three largest neurons in the HEMI-BRAIN ROI-1 demonstrated that UniSPAC-3D can acquire a satisfactory neuron tracing accuracy.

Online learning technology plays a crucial role in adapting segmentation models to new species. By leveraging UniSPAC-2D’s capability to segment and proofread vEM neurons, we can efficiently generate annotated data for previously unstudied species. Powered by the online learning strategy, UniSPAC-2D can quickly achieve better segmentation performance with the newly labeled data of new species. We simulated and validated this pipeline on Drosophila and zebra finch data using various online learning algorithms. After pre-training on Drosophila data and applying online learning on six volumes of zebra finch, the UniSPAC-2D model achieved increasingly better segmentation performance on the remaining zebra finch test volumes. Among the seven online learning algorithms chosen and compared here, the AGEM strategy delivered the best overall performance.

The key point of UniSPAC lies in its combination of 2D interactive segmentation and 3D tracking, distinguishing it from existing neuron segmentation methods. While its architecture shares some conceptual similarities with SAM2^43^ and FFN^23^, it introduces notable distinctions. In FFN, the process begins with an initial point input to generate the first output mask, which is then iteratively refined by re-inputting the output mask along with the raw vEM image, to achieve progressively better results. SAM2, on the other hand, segments the first slice (or frame) based on prompts (e.g., points or bounding boxes) and utilizes a memory bank mechanism to sequentially infer masks for subsequent frames, without reusing the previous output mask as input. In contrast, UniSPAC-3D builds upon UniSPAC-2D by segmenting the first slice using a prompt map to produce an initial 2D mask. This 2D mask is subsequently re-used as input for inferring masks across vEM blocks, enabling segmentation that extends beyond just the next immediate frame.

Despite the current process, several limitations need to be addressed in future work. First, while UniSPAC’s interactive features enable more accurate segmentation compared to approaches purely based on affinity graph^16,44^, they also require increased manual effort. Integrating automated proofreading tools, such as RoboEM^32^ and Gapr^45^, into UniSPAC could help identify potential segmentation errors and significantly enhance proofreading efficiency. Second, in the current application, UniSPAC utilizes vEM data from only two species. Expanding the dataset to include vEM data from various species could present a valuable opportunity—and a potential challenge—for training the foundation model for neuron segmentation in the future.

## Methods

### Datasets

To evaluate the UniSPAC models, we adopted vEM image data from two species for our experiments. The first species is the Drosophila melanogaster, which is one of the most thoroughly studied organisms. The other species is zebra finch, a popular model organism among neurobiologists interested in vocal learning. Data for both species come from a recent study^16^ for neuron segmentation. For Drosophila melanogaster, two datasets named HEMI-BRAIN^46^ and FIB-25^47^ from previous publications were combined to train neuron segmentation models. HEMI-BRAIN and FIB-25 were both imaged at 8 nm isotropic resolution using focused ion beam scanning electron microscope (FIB-SEM). HEMI-BRAIN has eight densely annotated volumes (∼450µm^3^), which took up 0.002% of the entire dataset. FIB-25 has four densely labeled volumes with ∼160µm^3^ of the original data. Following the recent study^16^, we adopted the HEMI-BRAIN ROI-1 (∼12µm edge length) to evaluate the performance of all approaches in segmenting neurons. For zebra finch, 33 densely labeled volumes were adopted as the datasets in cross-species segmentation. To validate the cross-species segmentation performance of all segmentation methods trained on Drosophila melanogaster data, all 33 zebra finch volumes were initially used as test data. Subsequently, the front 6 volumes were regarded as training data in online learning setup and were sequentially used to optimize the performance of the interactive segmentation model.

### UniSPAC-2D: an interactive segmentation model for 2D neuron images

In comparison with SAM, we proposed UniSPAC-2D for interactive neuron segmentation on 2D vEM slices. Two key differences in model architecture should be noted between SAM and UniSPAC-2D. First, SAM was primarily based on Vision Transformer (ViT), while UniSPAC-2D was constructed using multiple U-Net-based functional components. Second, the point prompts were regarded as tensors of coordinates and got embedded by positional encodings in SAM, whereas UniSPAC-2D converted all point prompts into prompt maps according to predefined rules. These above differences enhanced UniSPAC-2D’s accuracy and interactivity in neuron segmentation tasks. In general, UniSPAC-2D was composed of two functional modules: one for predicting neuron affinities and the other for predicting neuron masks. Motivated by ACRLSD, which was reported to be the state-of-the-art affinity-based method, we employed a 2D version of ACRLSD as the affinity prediction component of UniSPAC-2D. ACRLSD-2D processes raw 2D slices to predict neuron affinities. Besides, ACRLSD-2D also output local shape descriptors (LSDs), which were proposed as intermediate results to improve the quality of predicted affinity. The prediction for LSDs, affinity, and mask were implemented by three 2D U-Net models (Supplementary Figure 5a). For the first 2D U-Net network to predict LSDs, only one initial feature map representing the raw gray 2D image was adopted as input. The second 2D U-Net network to predict affinity utilized seven initial feature maps, including six channels for predicted 2D LSDs and one channel for raw gray scale images. These 2D U-Net networks constructed ACRLSD-2D, which was trained previously and got frozen during subsequent training for the mask prediction component of UniSPAC-2D. The final 2D U-Net network (mask prediction component) took the predicted affinity and the prompt map converted from point prompts according to predefined rules as input to acquire the final neuron mask.

In SAM^33^, the point and box prompts are represented by positional encodings^48^. A recent BioSAM^49^ model generates a superpixel-guided prompt to improve SAM’s segmentation performance. In our approach, UniSAPC-2D is built upon the U-Net model, renowned for a strong ability to capture local features through its effective receptive field. Therefore, we convert the point prompt of UniSAPC-2D into a map that aligns with the dimensions of the input image. The prompt map (PM) was generated from point prompts according to predetermined rules:

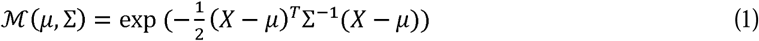

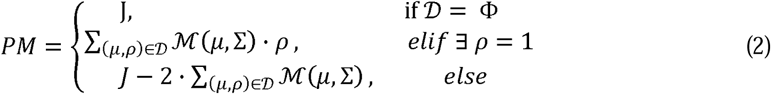

where D is the set of point prompts, μ is the position of one point prompt and ρ ∈ {-1,1} is its type, 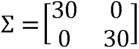 is the covariance matrix, *X* represents all pixels on the 2D image, J is a full one matrix that has same shape like the 2D image. We designed the PM in this way since there exists three situations. First, we hope that UniSPAC-2D can still perform segmentation even when no prompts are given. Therefore, we set the PM to default values (l) and UniSPAC-2D is supposed to predict masks for all neurons. Second, when positive prompts (ρ = 1) are given, UniSPAC-2D should segment only the neurons chosen by positive prompts, delete areas with negative prompts (ρ = -1) and neglect other areas with no prompts. Third, when only negative prompts are given, UniSPAC-2D need to segment all neurons that are not selected by negative prompts.

### UniSPAC-3D: a 3D neuron tracking model based on UniSPAC-2D

UniSPAC-3D can segment selected neurons across slices by the prompts from UniSPAC-2D, exponentially reducing the amount of interaction workload. The architecture of UniSPAC-3D comprised three modules: one affinity module to predict the 3D affinity map, the UniSPAC-2D module to predict the 2D mask of neurons in the first slice, and the final mask module to predict the 3D mask of selected neurons. Despite UniSPAC-2D, the other two modules in UniSPAC-3D were both implemented using 3D U-Net networks (Supplementary Figure 5b). In line with the design of UniSPAC-2D, we adopted the 3D version of ACRLSD as the module to predict the affinity map in UniSPAC-3D. Then, the first slice in the 3D raw block was employed as an input of UniSPAC-2D that offers an interactive 2D segmentation. Furthermore, the resulting 2D masks of the first slice would replace the raw first slice in the 3D raw block. UniSPAC-3D finally utilized a 3D U-Net network to predict the 3D masks of selected neurons from the spliced 3D affinity map and the updated 3D raw block. All 3D U-Net networks in UniSPAC-3D were down-sampled by a factor of [2, 2, 2] in each layer. The output logits for 3D masks in the last 3D U-Net were constrained between 0 and 1 through a sigmoid function.

### The regularization term of the loss function

In order to enhance the continuity of neuronal boundaries within the masks predicted by the UniSPAC-2D model, a loss regularization term has been devised:

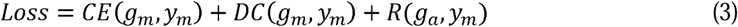

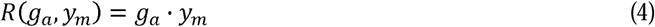

where *CE*(,)refers the binary cross entropy loss function, *DC*(,) refers the dice loss function, *R*(,) refers the loss regularization term, *g_a_* ∈ {0,1} is the ground truth mask for a voxel, *y_m_*is the prediction for its mask, and *g_a_* ∈ {0,1} is the ground truth affinity for the voxel. For those voxels that are on the boundary (*g_a_* = 0,1), their *y_m_* should be as close to 0 as possible.

### Training

Both UniSPAC-2D and UniSPAC-3D models were composed of multiple functional modules, and each module was trained independently. Two ACRLSD models adopted in UniSPAC-2D and UniSPAC-3D were both trained with a primary loss targeting affinity and an auxiliary loss for LSDs. Following the previous study^16^, the loss function for ACRLSD models was based on the mean squared error (MSE), and minimized using the Adam optimizer. For the training of UniSPAC-2D and UniSPAC-3D models, a combination of cross entropy loss and dice loss was adopted. In the cross-species segmentation experiment, we also introduced an enhanced version of UniSPAC-2D, named UniSPAC-2D*. UniSPAC-2D* further required a regularization term beyond the combined loss during training and significantly improved the quality of predicted neuronal boundaries.

Importantly, ACRLSD models were frozen during the training for UniSPAC-2D and UniSPAC-3D models. Each densely annotated sample was loaded in one of two modes: non-prompt or with-prompt, with a 50% probability assigned to each mode. In non-prompt mode, all labeled neurons should be segmented and the *PM* should be set to default value of J since no point prompts were provided. In with-prompt mode, only neurons selected by the point prompts should be segmented and the *PM* should be established according to predefined rules. Generally, the ground truth mask of the UniSPAC models was the mask of all neurons when the prompt was absent or the mask of selected neurons when the prompt was provided. If an vEM image sample was loaded in with-prompt mode, we randomly selected the neurons to be segmented with a probability ranging from 0.1 to 0.9. The details of the above data loading process were consistent for both UniSPAC-2D and UniSPAC-3D. In addition, the ground truth of masks for the first slice of the 3D raw block was directly adopted to replace the original first raw slice during the training of UniSPAC-3D. All training processes were performed under an early-stop strategy which terminates training if its loss on the validation set does not decrease for 100 consecutive epochs. The training process was performed on multiple 40G-A100 GPUs, with the learning rate set to 0.0001. All 2D networks took an input shape of [128, 128], and all 3D networks accepted an input shape of [32, 128, 128], except that the validation input shape for 3D networks was set to [8, 128, 128].

### Methodology of online learning

Online Learning (OL) facilitates the continuous adaptation to new tasks for existing segmentation models while retaining previously acquired knowledge. Typically, OL approaches are categorized based on their strategies for addressing the forgetting problem: 1) regularization-based; 2) replay-based; 3) distillation-based^38^. This section outlines six key OL methods: Replay, Cumulative Learning, Elastic Weight Consolidation (EWC), Synaptic Intelligence (SI), GDumb, and Averaged Gradient Episodic Memory (A-GEM).

Replay methods mitigate catastrophic forgetting by revisiting stored past experiences during training on new tasks, thereby preserving prior data distributions. Cumulative Learning involves retraining the model on the entire dataset, including both old and new data. This approach ensures the retention of all past knowledge, allowing the model to benefit from a comprehensive history of experiences. Elastic Weight Consolidation (EWC)^39^ protects previously learned knowledge by incorporating a regularization term into the loss function. This term penalizes changes in weights considered important for earlier tasks, balancing the acquisition of new knowledge with the preservation of existing information, and enabling the model to adapt to new tasks without significant loss of previously learned skills. Synaptic Intelligence (SI)^40^ extends the principles of EWC by dynamically estimating the importance of each parameter based on its contribution to loss reduction. SI adjusts learning rates to protect significant parameters, making it particularly suitable for task-incremental learning scenarios. GDumb (Greedy Sampler and Dumb Learner)^41^ utilizes a straightforward strategy of storing samples greedily and training a model from scratch using these stored samples at test time. This approach avoids complex mechanisms and focuses on maintaining balanced class representation in memory, resulting in competitive performance across various CL scenarios. Averaged Gradient Episodic Memory (A-GEM)^42^ is a gradient-based memory method that prevents forgetting by adjusting the gradient during training. Unlike GEM, which uses multiple constraints, A-GEM simplifies the process by employing a single average constraint. This approach ensures that the current gradient direction does not increase the loss on past tasks, thereby maintaining stability while learning new tasks.

These methods encompass a variety of strategies for OL, each employing distinct mechanisms to address catastrophic forgetting problem. Replay and Cumulative Learning mainly focus on comprehensive data retention. EWC and SI prioritize the preservation of crucial parameters. A-GEM offers a streamlined gradient-based solution. GDumb exemplifies the effectiveness of a minimalist design. We utilized 33 volumes of zebra finch data to evaluate the performance of these OL methods. The first six volumes were designated as the training data stream, while the remaining 27 volumes served as the testing data. These OL strategies were compared on both segmentation accuracy and training time to identify the most effective strategy for practical application.

## Data availability

All datasets adopted in this study are publicly available at https://github.com/ddd9898/UniSPAC/tree/main/data along with a detailed tutorial for reproducing the reported results. A demo of 2D interactive segmentation and 3D tracking using UniSPAC can be found in Supplementary Video 1.

## Code availability

A PyQt-based implementation of the UniSPAC tool can be found at our GitHub repository: https://github.com/ddd9898/UniSPAC/tree/main. All source code to train and evaluate UniSPAC-2D and UniSPAC-3D models are also publicly available with the MIT license.

## Acknowledgements

This work is Supported by Shanghai Artificial Intelligence Laboratory.

## Author contributions

C.S., W,O., J.P.W. and J.D conceived this project. C.S., W.O. and J.P.W. supervised this research. J.D., J.M.W. and

C.S. designed detailed implementations for this study and J.D. conducted the experiments. C.C. completed the replication of the online learning algorithms and the writing of the related section. J.M.W. and Q.Z. helped to perform data analysis, participated in critical discussions about the results and contributed to the revision of the manuscript. Z.Z. and J.P.W. provided significant comments and guidance on the experiment design and participated in the revision of the paper. C.S. and W.O. provided critical support on the system setup, including the computing platform. All authors discussed the results and commented on the manuscript.

