## Supplementary Information for "UniSPAC: A Unified Segmentation Framework for Proofreading and Annotation in Connectomics"

- **Supplementary Note 1: Testing of the patch size parameter in UniSPAC-3D.**
- **Supplementary Figure 1: Schematic of UniSPAC-3D chunking inputs in the XY plane while tracking HEMI-BRAIN-ROI-1.**
- **Supplementary Table 1: Comparison of different patch size settings for UniSPAC-3D tracking.**
- **Supplementary Figure 2: Visualization of neuron tracking results with different patch size parameters.**
- **Supplementary Figure 3: Comparison of VOI split metrics for 33 zebra finch volumes by different methods in the cross-species segmentation experiment.**
- **Supplementary Figure 4: Comparison of VOI merge metrics for 33 zebra finch volumes by different methods in the cross-species segmentation experiment.**
- **Supplementary Table 2: Comparison of inference elapsed time between UniSPAC-2D and ACRLSD-2D on the same hardware.**
- **Supplementary Figure 5: Schematic representation of the structure and parameters of each U-Net network in UniSPAC-2D and UniSPAC-3D.**
- **Supplementary Video 1: A video showing how UniSPAC interactively segments neurons and proofreads for errors.**

**Supplementary Note 1: Testing of the patch size parameter in UniSPAC-3D.**

The patch size parameter of UniSPAC-3D controls the number of slices in which neurons are automatically tracked at each inference. The patch size must be a multiple of 8 since the down-sampling factors of 3D U-Net in UniSPAC-3D were [[2,2,2], [2,2,2], [2,2,2]], [2,2,2]]. We tested different patch sizes (8, 16, 24, 32) to explore the limits of UniSPAC-3D's auto-tracking capabilities. For HEMI-BRAIN-ROI-1 (1475×1475×1475), when the patch size is set to 8, UniSPAC-3D processes a raw image block of 8×1480×1480 for each inference. However, when the patch size is set to 16, 24, or 32, the GPU memory (40GB) imposes limitations, preventing inference on such large raw image blocks. To overcome this, we divided the image into smaller chunks along the XY plane (Supplementary Figure 1). The size of each chunk in the XY plane is 512×512, resulting in 9 chunks in the XY plane. Thus, for patch sizes of 16, 24, or 32, the final input to UniSPAC-3D becomes 16×512×512, 24×512×512, or 32×512×512, respectively.

For the three neurons with the largest volumes in the HEMI-BRAIN-ROI-1, we adopted UniSPAC-3D to trace them across slices under different patch sizes (8, 16, 24, 32). As shown in Supplementary Table 1, the tracking accuracy (VOI) of UniSPAC-3D for these three neurons in HEMI-BRAIN-ROI-1 gets worse as the patch size increases. When the patch size is set to 32, significant noise appears in the tracking results for neurons 57, 68, and 184. In addition, the neurons are not reconstructed with sufficient detail. In practice, in order to balance precision and efficiency, we still recommend using a patch size of 8 or 16.

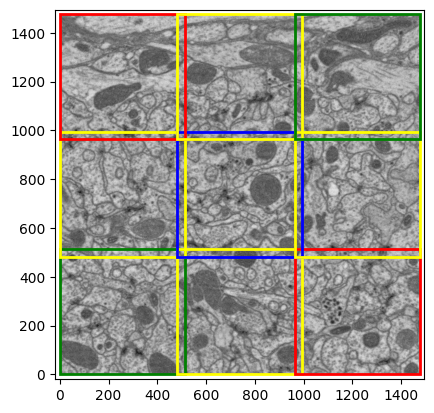

**Supplementary Figure 1: Schematic of UniSPAC-3D chunking inputs in the XY plane while tracking HEMI-BRAIN-ROI-1.**

**Supplementary Table 1: Comparison of different patch size settings for UniSPAC-3D tracking.**

| **Patch Size** | **VOI (Neuron 68)** | **VOI (Neuron 184)** | **VOI (Neuron 57)** |
| --- | --- | --- | --- |
| **8** | 0.061 | 0.075 | 0.087 |
| **16** | 0.100 | 0.110 | 0.161 |
| **24** | 0.127 | 0.154 | 0.232 |
| **32** | 0.140 | 0.266 | 0.267 |

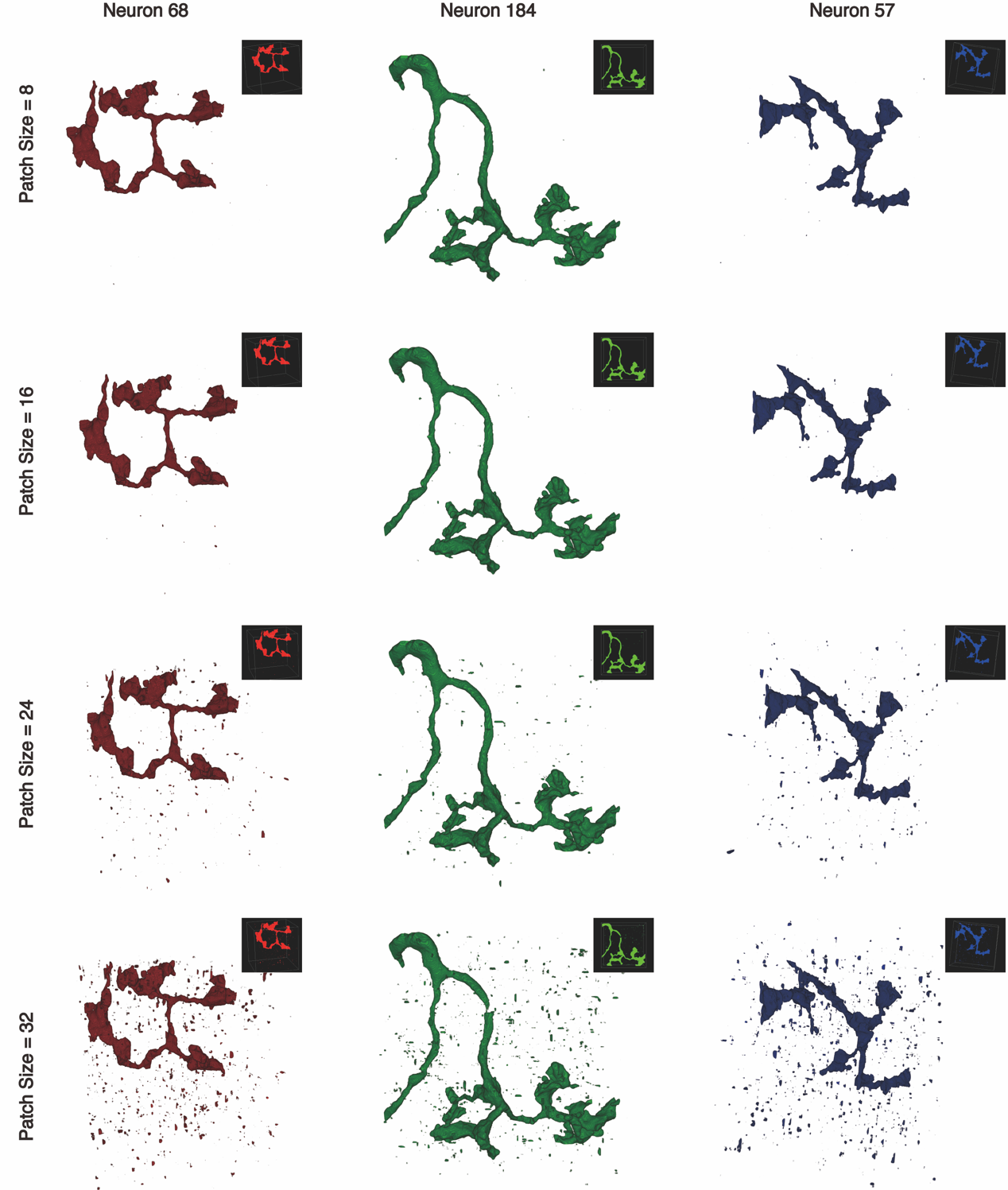

**Supplementary Figure 2:** **Visualization of neuron tracking results with different patch size parameters.**

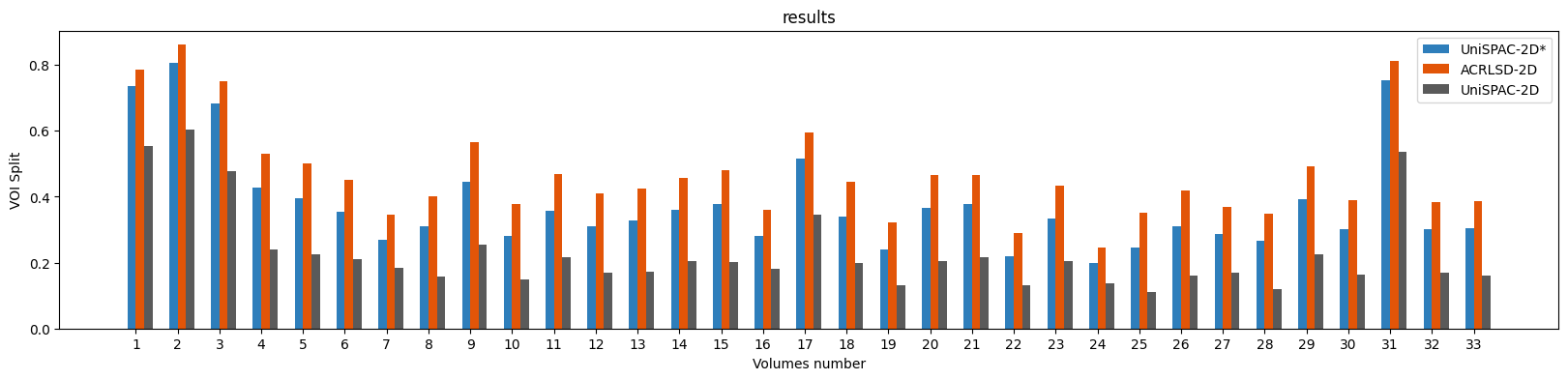

**Supplementary Figure 3: Comparison of VOI split metrics for 33 zebra finch volumes by different methods in the cross-species segmentation experiment.**

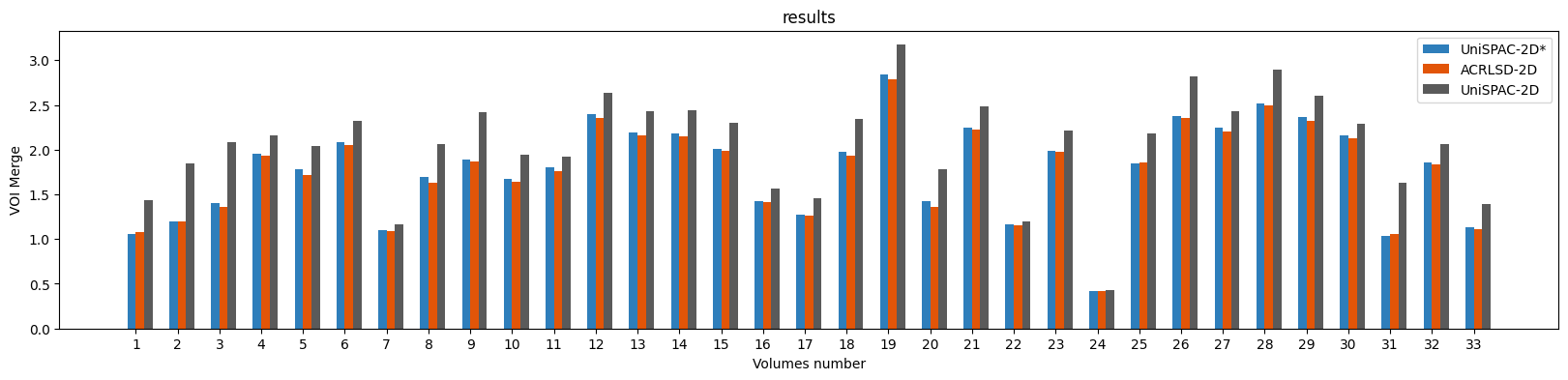

**Supplementary Figure 4: Comparison of VOI merge metrics for 33 zebra finch volumes by different methods in the cross-species segmentation experiment.**

**Supplementary Table 2: Comparison of inference elapsed time between UniSPAC-2D and ACRLSD-2D on the same hardware.**

|  | **UniSPAC-2D** | | | **ACRLSD-2D** | | |
| --- | --- | --- | --- | --- | --- | --- |
| **Image size** | **GPU time** | **CPU time** | **Total time** | **GPU time** | **CPU time** | **Total time** |
| 256*256 | 0.0322 | 0.0008 | **0.0330** | 0.0051 | 0.0445 | 0.0496 |
| 512*512 | 0.0268 | 0.0027 | **0.0295** | 0.0063 | 0.1490 | 0.1553 |
| 1024*1024 | 0.0320 | 0.0098 | **0.0418** | 0.0052 | 0.7233 | 0.7285 |
| 2048*2048 | 0.0566 | 0.0397 | **0.0963** | 0.0052 | 3.6788 | 3.6840 |

* Time units are in seconds. The GPU is one A100 with the 40 GB memory.

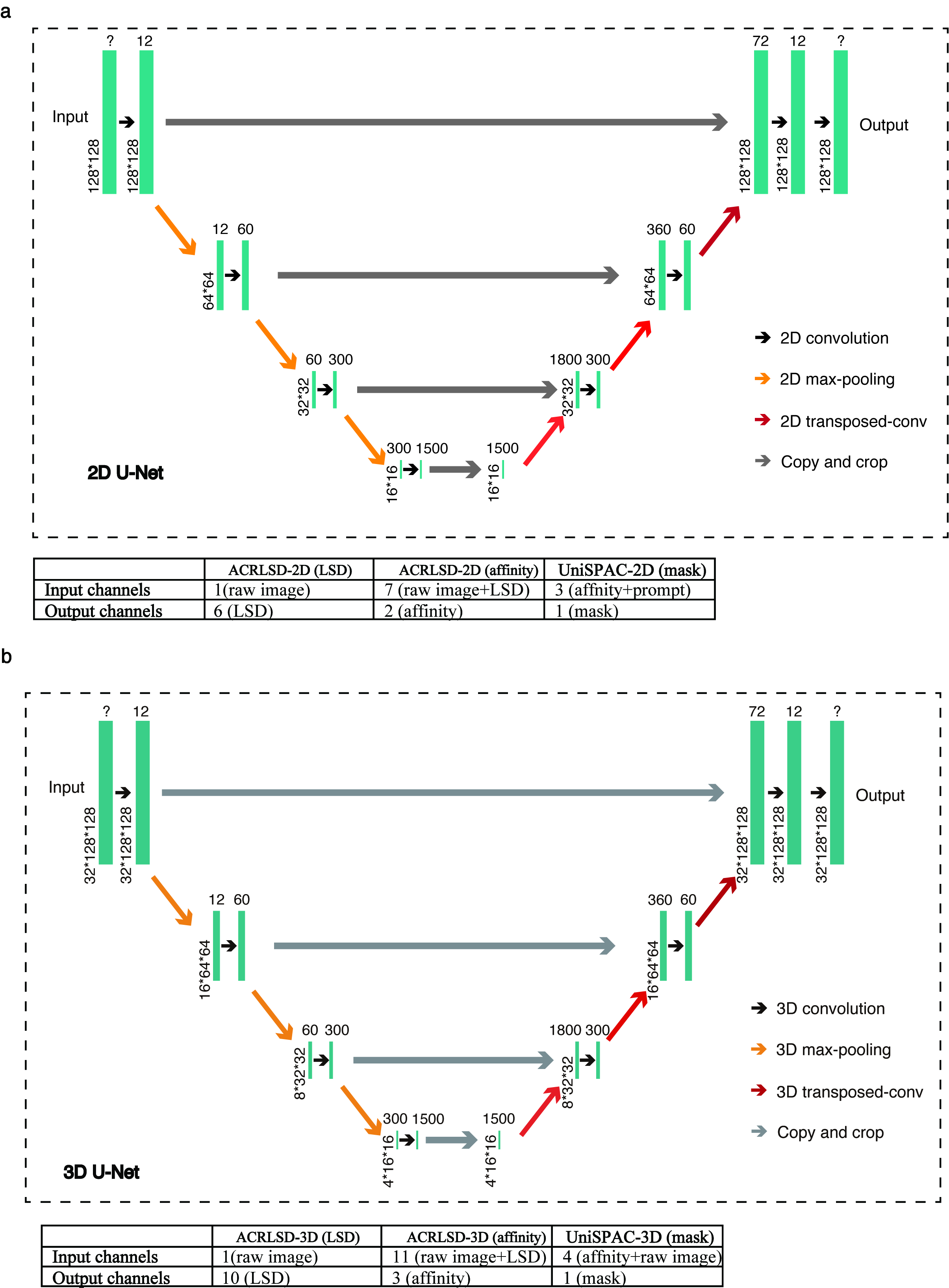

**Supplementary Figure 5: Schematic representation of the structure and parameters of each U-Net network in UniSPAC-2D and UniSPAC-3D.**
